# TACO: An open, interpretable model for microsecond-scale TCR specificity prediction

**DOI:** 10.64898/2026.11.04.764339

**Authors:** Marco Garcia Noceda, Erin Calfee, Brian C. Zhang, Edward Osborne, Brad Greenfield, Mark Klinger, Matthew T. Noakes, Bryan Howie, Harlan Robins

## Abstract

Predicting antigen-induced T-cell activation from T-cell receptor (TCR) and peptide sequence remains difficult for TCRs and peptides not represented in training data. We present TACO (T-Cell Activation from Chemistry and Orientation), an open statistical model for predicting TCR specificity against HLA-A*02:01-presented peptides. TACO takes as input the peptide sequence and six TCR CDR loop sequences. Learning from binary activation labels, TACO selects a single canonical orientation of the CDR loops over the peptide and learns five amino-acid contact-potential matrices as well as two peptide-exposure profiles. The orientation is fixed and does not require experimentally determined or predicted structures. We evaluate TACO on held-out peptides separated by Levenshtein distance at least four from all training and validation peptides, limiting near-neighbor shortcut learning. TACO achieves AUROC 0.67 on a large held-out activation dataset and 0.68 on IMMREP25 HLA-A*02:01, exceeding the open structure-based models evaluated here and matching the reported AlphaFold3-TCRdock result. The model has 2,019 fitted parameters, retains most within-peptide ranking signal with TCR*β* sequence alone, and reaches 406,000 TCR–peptide predictions per second on CPU. This throughput enables computational evaluation of large TCR–peptide search spaces. We show its potential use for prioritizing candidate target peptides from large peptide libraries, reducing the search space for downstream experimental investigation. TACO fitted parameters also expose candidate physicochemical patterns that can be inspected directly. We release TACO’s parameters and complete training, evaluation, and inference code.

## 1 Introduction

T cells recognize peptides presented by major histocompatibility complex (MHC) molecules via their T-cell receptor (TCR). In humans, MHC molecules are called human leukocyte antigen (HLA) molecules. Predicting whether a T cell expressing a particular TCR will activate in response to a peptide presented by a given class I HLA molecule is an important and challenging problem in computational immunology [10, 19, 7]. The vast combinatorial space of peptide sequences [23], class I HLAs [14], and TCRs [12] makes dense experimental coverage of TCR–pMHC interactions intractable. Available activation data cover only a small fraction of these combinations [24][21], and experimentally determined structures of TCRs bound to peptide-MHC (pMHC) complexes are scarcer still [13]. Computational methods are needed to extend these limited experimental observations to previously untested TCR-pMHC combinations.

Such computational methods, provided they achieve useful generalization to peptides dissimilar to their training data, would ideally also support large screens of many TCRs against pMHCs at low computational cost and provide insight into how individual interactions within the TCR-pMHC complex contribute to their predictions. Recent benchmarking has demonstrated that AlphaFold3-based pipelines have leading predictive signal on novel peptides [19]. However, these pipelines are computationally expensive, requiring a predicted structure for each TCR-pMHC pair. Additionally, although their predicted structures can be examined, these pipelines infer specificity from confidence measures of the structural prediction or from heuristic scores of its modeled interface, and these scores do not directly reveal the interactions responsible for functional activation. Deep sequence models have been shown to achieve competitive performance and faster inference when trained on sufficiently large activation datasets [17]. However, individual interaction contributions remain difficult to inspect, and repertoire-scale screens requiring billions of predictions can still become expensive for these models.

Interaction-oriented models, including TCRen [9] and TCRcube [5], provide starting points for representing recognition through residue interactions. Introspection of the deep sequence model ImmSET further suggested that simple electrostatic features capture part of its predictive signal [17]. These findings motivate research into a complementary strategy: using structural priors and hypotheses suggested by expressive sequence models to construct compact statistical predictors with explicit interaction terms.

Here, we introduce TACO (T-cell Activation from Chemistry and Orientation), an open statistical model for predicting antigen-induced T-cell activation by HLA-A*02:01-presented peptides. Training on the large activation dataset from the ImmSET study, TACO learns amino-acid pair scores and peptide-position weights. A shared approximation of how receptor binding loops sit over the peptide allows predictions without modeling a separate structure for each TCR–peptide pair. On HLA-A*02:01 peptides separated from all training and validation peptides in sequence space, TACO matches the reported performance of a leading AlphaFold3-based pipeline and exceeds the open structure-based models evaluated in this study. With only 2,019 fitted parameters, the model supports CPU inference at up to 400,000 TCR–peptide predictions per second, and exposes candidate interaction patterns involving electrostatic complementarity, charge-selective interactions consistent with structural mimicry, and sulfur–aromatic chemistry. We release the fitted parameters and complete training, evaluation, and inference code.

Additionally, we conduct a scaling analysis to evaluate how TACO improves as a function of training data and to contextualize its performance relative to higher-capacity neural-network-based models. TACO is competitive with individual high-capacity models in smaller data regimes, but its performance gains flatten over the range studied while less restricted sequence models continue to benefit from additional data. This comparison does not identify a universal best model class, but suggests complementary roles: large activation datasets and high-capacity models can uncover predictive signal, while compact models such as TACO can make part of that signal accessible for hypothesis generation and experimental follow-up.

## 2 The TACO Model

TACO’s inputs are the peptide sequence and the six CDR-loop sequences of the TCR carried by that T cell, and its output is the estimated probability of the positive binary activation label. The model is trained against binary labels of functional T cell activation generated by the MIRA (Multiplexed Identification of T cell Receptor Antigen specificity) assay [11]. It operates directly on sequence and does not predict or co-fold a TCR–pMHC structure for each TCR–peptide pair, instead using a low-dimensional canonical orientation selected from a small set of candidate orientations using validation data. This canonical orientation represents the position of the TCR over the peptide [6] and defines where each CDR loop can contribute. Within this canonical orientation, TACO learns amino-acid interaction preferences and peptide-position weights from the activation labels.

Figure 1 summarizes the model and its fitted components. The six loops contribute through five loop groups: CDR1 *α*, CDR2 *α*, CDR1*β*, and CDR2*β* each define one group, while CDR3 *α* and CDR3*β* share a fifth group. A loop group is a set of one or two loops that share a contact-potential matrix. We refer to the four CDR1/2 loop groups as germline loop groups because these loops are encoded by the TCR variable gene segments.

**Fig. 1:**
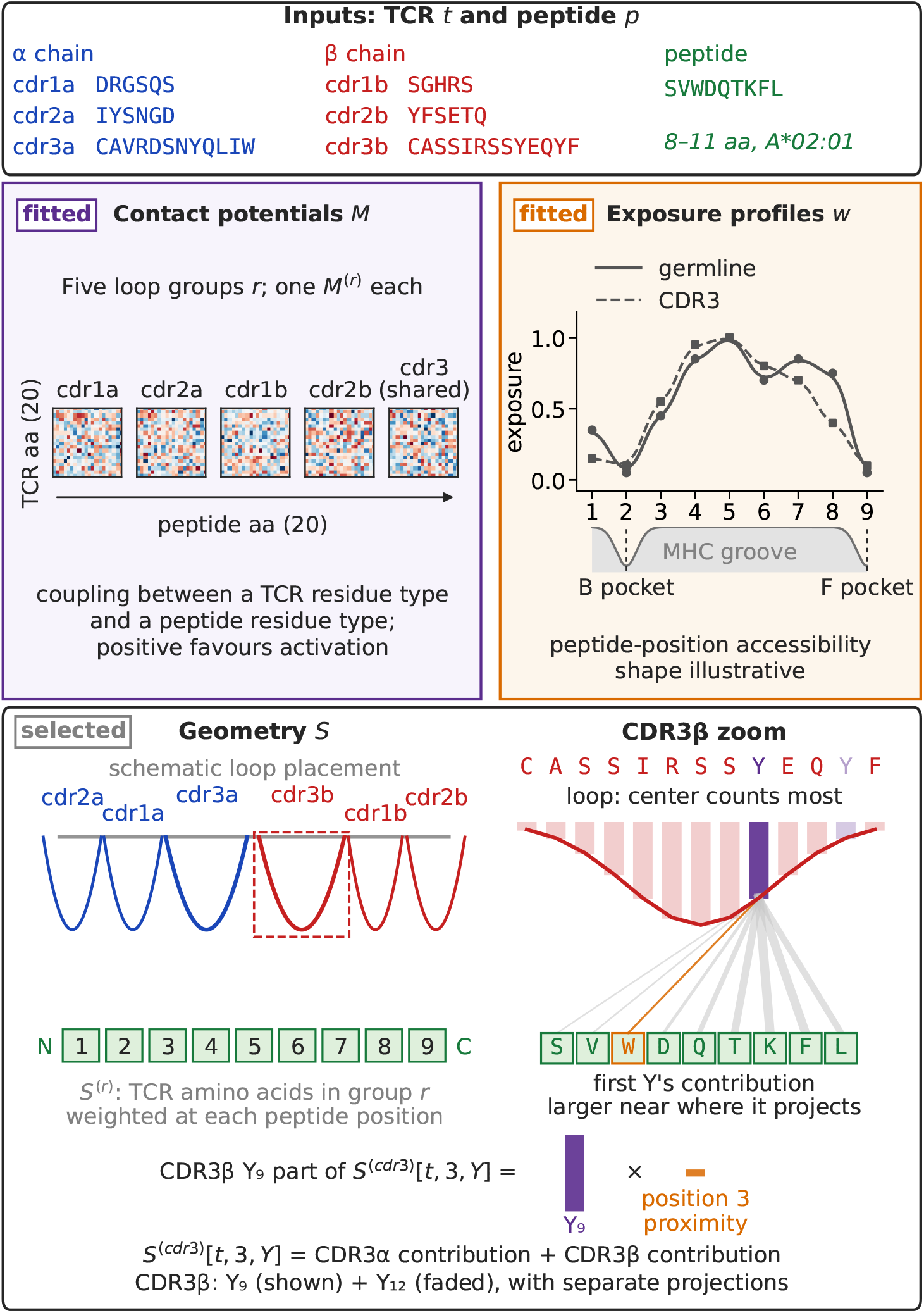
Overview of TACO. Schematic geometry weights each TCR loop residue by its position within the loop and its proximity to peptide positions; the CDR3*β* zoom illustrates one contribution. Five learned contact-potential matrices score amino-acid pairs, while two learned exposure profiles down-weight peptide positions buried in HLA-A*02:01.

### Contact potentials

Each loop group *r* has an independent 20 × 20 contact-potential matrix, *M* ^(*r* )^ [*m, n*], whose rows index amino-acid types in the TCR loops and whose columns index amino-acid types in the peptide. The entry *M* ^(*r*)^ [*m, n*] gives the learned contribution when TCR residue type *m* is presented to peptide residue type *n* through loop group *r*. The matrices are fitted directly from the binary activation labels and are otherwise unconstrained. Thus, each loop group learns a distinct, directionally defined map of TCR–peptide residue interactions. Retrospective comparisons testing different model variants favored five contact-potential matrices over versions with fewer separate loop groups, and found no improvement from adding in separate CDR3 *α* and CDR3*β* potentials (Appendix A).

### Peptide exposure

Peptide positions differ in their accessibility to the TCR: in HLA-A*02:01, residues at the B and F anchor positions are commonly buried in the MHC groove, whereas central residues are generally more exposed [20]. TACO represents this variation with learned exposure profiles *w*_*j*_, which scale contact contributions at peptide position *j*. One profile is used for the four germline loop groups and one for the shared CDR3 loop group. Each profile is specified by nine control points and interpolated to peptide length. The two-profile variant performed better than a shared profile (Appendix A).

### Orientation

An orientation specifies, for each loop *l*, a peptide-coordinate center *µ*_*l*_, sweep slope *ρ*_*l*_, and contact width *σ*_*l*_. The center *µ*_*l*_ describes the peptide position with which loop *l* interacts most strongly. For example, *µ*_*l*_ = 5 places the center of loop *l* over position 5 of a 9-mer peptide. The sweep slope *ρ*_*l*_ specifies how the projected position of the loop changes across residues of the loop. The contact width *σ*_*l*_ specifies the breadth of the loop footprint over the peptide: larger values allow a loop residue to contribute to a wider range of nearby peptide positions, whereas smaller values concentrate its contribution near its projected position.

We evaluated seven hand-specified orientations that varied these parameters (Appendix B). For each orientation, the contact potentials and exposure profiles were fitted independently, and the canonical orientation was selected by validation AUROC. The selected orientation retains the hand-specified germline-loop spacing, places the CDR3 *α* and CDR3*β* centers two peptide-coordinate units to either side of the peptide midpoint, uses contact widths of2.0and2.6for germline and CDR3 loops, respectively, and has zero sweep slope. Consequently, all residues in a loop share a projected center over the peptide, with the Gaussian coupling below giving the loop a footprint spanning multiple peptide positions.

The geometry first assigns each residue a weight according to its position within its loop. For loop *l* of length

*L*_*l*_,let

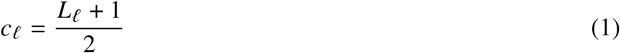

denote the midpoint of the loop. Residue *i* receives within-loop weight

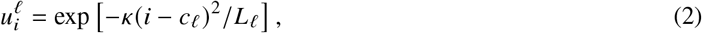

Where *κ* is fixed. Thus, residues near the midpoint of the loop contribute more strongly than residues near either end.

For a peptide of length *L*_*p*_, residue *i* in loop *l* projects to peptide coordinate

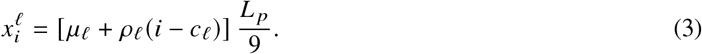

Its coupling to peptide position *j* is

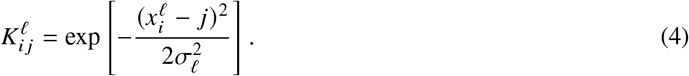

Thus,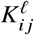 is largest when loop residue *i* projects near peptide position *j* and decreases smoothly with distance.

For each TCR *t*, loop group *r*, peptide position *j*, and TCR amino-acid type *m*, loop residues are then aggregated into the intermediate feature

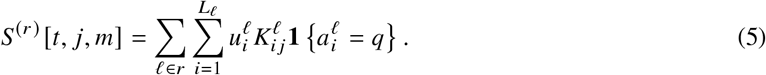

Here,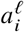 is the amino acid at loop residue *i*. Thus, *S* ^(*r*)^ [*t,j,m*] records the geometry-weighted amount of aminoacid type *m* from loop group *r* presented to peptide position *j*. It is determined by the TCR loop sequences and lengths, the selected orientation, and peptide length; it does not depend on peptide amino-acid identities.

### Contact score and activation probability

The overall contact score sums these intermediate TCR features across loop groups and peptide positions. At each position, *S* ^(*r*)^ [*t, j,m* ] specifies how strongly a TCR amino-acid type is presented, *M* ^(*r*)^ [*m, n*_*j*_] specifies the learned contribution of pairing it with the peptide amino acid, and the exposure profile scales that contribution according to peptide accessibility. For TCR *t* and peptide *p*, the contact score is

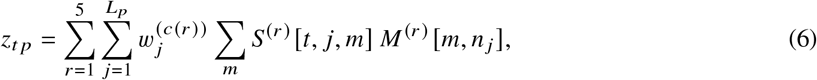

Where *n*_*j*_ is the peptide amino acid at position *j* and *c* (*r*) assigns loop group *r* to the germline or CDR3 exposure profile. The predicted probability of the positive activation label is

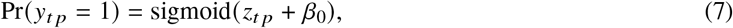

where *β*_0_ is a global intercept.

### Fitting

For each candidate orientation, TACO learns five contact-potential matrices, two peptide-exposure profiles, and a global intercept from the observed TCR–peptide activation labels. Parameters are fitted by minimizing binary cross-entropy over all training rows, with a small ridge penalty (*λ* =10 ^−6^) on the contact potentials. Fitting alternates between two parameter updates. With the exposure profiles held fixed, TACO optimizes the contact-potential matrices and global intercept; with the contact potentials held fixed, it optimizes the exposure profiles. Each update uses the full training set without class rebalancing, thereby preserving the observed class imbalance. Optimization and implementation details are provided in Appendix C.

## 3 Evaluation

### 3.1 Evaluation sets and peptide-distance holdout

The main TACO model is fitted on Dataset 0, the HLA-A*02:01 training set introduced previously [17]. This dataset contains 500 peptides, with 180 TCRs observed to activate in response to each peptide, together with the associated negative TCR–peptide examples. The labels in these datasets reflect antigen-induced T-cell activation following peptide stimulation, rather than a direct measurement of TCR–pMHC binding affinity; accordingly, TACO is evaluated here as a predictor of functional activation. For the scaling analysis, we reuse Datasets 0, 1, and 2 from [17], together with their predefined nested subsets from that study. All training, validation, and held-out partitions are therefore inherited unchanged.

We evaluate the final model on the primary held-out set and on the HLA-A*02:01 subset of IMMREP25. In both evaluations, every test peptide has Levenshtein distance at least four from every peptide in the training and validation sets. This criterion addresses a shortcut-learning failure mode in TCR–peptide prediction: nearby peptides can share TCR recognition patterns, allowing TCR–label associations within a peptide neighbourhood to produce apparent test-set signal without peptide-dependent recognition. The holdout therefore evaluates generalization to sequence-distant peptides.

### 3.2 Comparison with structural models

We estimate 95% confidence intervals by resampling peptides with replacement and recomputing AUROC. This peptide-level bootstrap captures variation in predictive performance across peptide targets rather than treating TCR–peptide rows as independent observations.

On the primary held-out activation set, which contains 134 HLA-A*02:01-restricted peptides with 50 positive and 200 negative TCRs per peptide, TACO obtains an AUROC of 0.67 [0.65, 0.69]. On the full 5,000-row HLA-A*02:01 IMMREP25 dataset, using all six CDR loops from the TCR *α* and TCR*β* chains, TACO obtains an AUROC of 0.68 [0.63, 0.73].

Table 1 compares TACO with the public structural models OpenDDE and OpenFold3 “preview 2” (OF3p2), which we evaluated in this study (Appendix D). The AF3-TCRdock metric was reported previously in [17]; we did not run AF3-TCRdock or generate predictions with it.

**Table 1:** Performance on the full 5,000-row HLA-A*02:01 IMMREP25 dataset. This analysis uses all released HLA-A*02:01 rows rather than the IMMREP25 competition evaluation subset. Confidence intervals for models evaluated in this study are estimated by peptide-level bootstrap resampling. The AF3-TCRdock value is taken from [17].

| Model | AUROC [95% CI] | Metric source |
|---|---|---|
| TACO | 0.68 [0.63, 0.73] | This study |
| OpenDDE | 0.63 [0.54, 0.71] | This study |
| OF3p2 | 0.58 [0.50, 0.66] | This study |
| AF3-TCRdock | 0.66 [–] | Reported in [17] |

**Table 2:** CPU inference throughput for TACO. “Panel mode” evaluates each TCR against multiple peptides and reuses shared intermediate quantities.

| Scenario | Evaluations | Single<br>time (s) | Single<br>pred./s | Multi<br>time (s) | Multi<br>pred./s |
|---|---|---|---|---|---|
| 10K TCRs × 1 peptide | 10,000 | 0.31 | 31,951 | 1.61 | 6,224 |
| 10K TCRs × 100 peptides | 1,000,000 | 8.71 | 114,839 | 9.16 | 109,182 |
| 1M TCRs × 1 peptide | 1,000,000 | 55.67 | 17,964 | 26.53 | 37,692 |
| 1M TCRs × 100 peptides | 100,000,000 | 414.48 | 241,267 | 246.21 | 406,159 |

TACO is trained specifically for HLA-A*02:01 and does not transfer comparably to other alleles. Across large held-out sets from additional HLAs, TACO retains non-null but modest predictive signal, including for HLA-B*40:01, but its performance remains substantially below that achieved in HLA-A*02:01 (Appendix E). This limited cross-HLA generalization is an important limitation of the current model. Further work is needed to determine whether similarly large allele-specific datasets can support comparable gains for other HLAs, and whether TACO-style models can eventually learn shared representations that generalize across multiple alleles.

### 3.3 Contribution of the TCR *α* chain

Because full repertoire profiling often sequences only the more diverse TCR*β* chain, we evaluated whether TACO retains predictive signal when TCR *α* sequence information is unavailable. For this analysis, we removed all TCR *α* loop contributions when constructing TACO’s geometry-derived features at inference time. The resulting score therefore uses only CDR1*β*, CDR2*β*, and CDR3*β* contributions; the learned contact potentials and peptide-exposure profiles are otherwise unchanged.

This approach directly evaluates the information provided by TCR*β* without introducing a synthetic TCR*α* sequence or requiring a peptide-specific score correction. On the IMMREP25 A*02:01 subset, the TCR*β*-only model obtains an AUROC of 0.65 [0.61, 0.70], a decrease of only 0.03 relative to the full-TCR model. Thus, TACO retains substantial predictive signal using TCR*β* alone, supporting applications to unpaired repertoire data.

## 4 Scaling Analysis

We assessed how TACO responds to additional training data using the predefined nested subsets of Datasets 0–2 [17]. For each fixed number of positive TCRs retained per peptide, we varied the number of training peptides and evaluated mean per-peptide AUROC.

TACO improves most rapidly at the smallest training-set sizes, with diminishing gains as additional peptides are included (Figure 2 for 180 TCRs/peptide result, full result in Appendix F). The fitted power-law curves do not fully describe the observed behaviour: performance flattens over the range examined. This pattern is consistent with a compact model whose contact potentials, exposure profiles, and orientation become better estimated with more data, but whose representational capacity remains fixed.

**Fig. 2:**
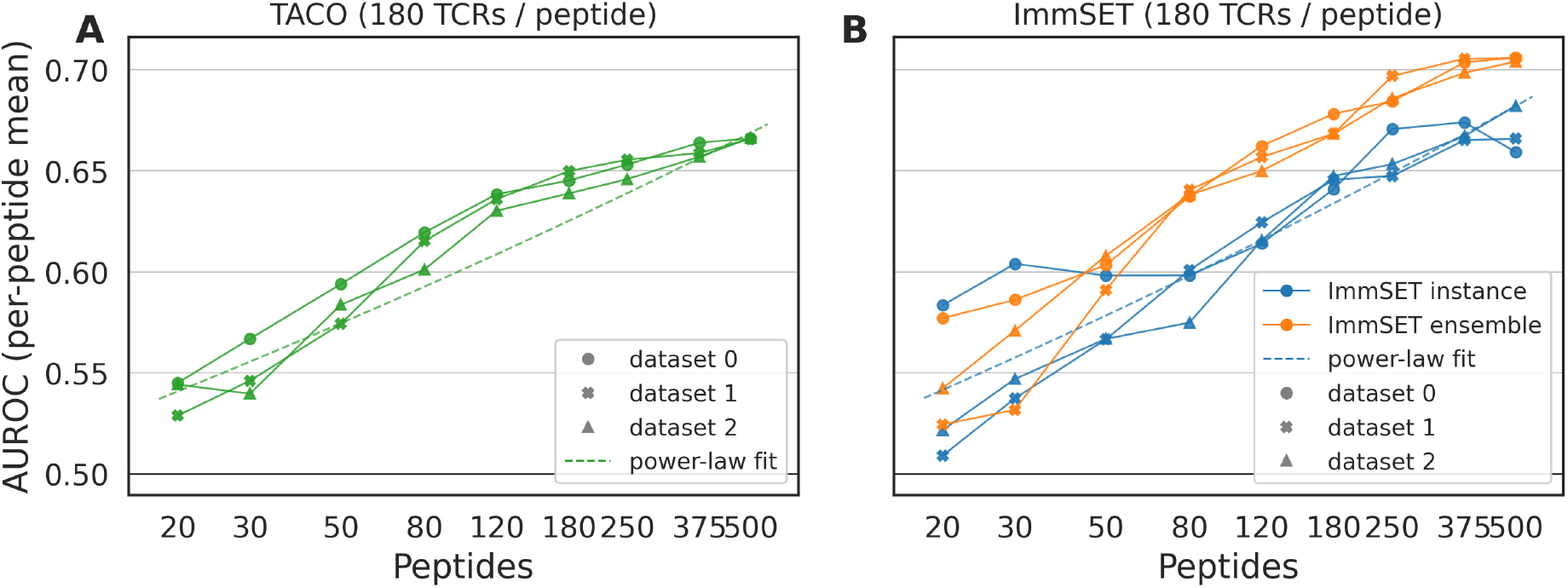
Scaling of TACO and ImmSET with training-data size. (A) TACO improves as the number of training peptides increases, but the gains diminish at larger training-set sizes; its observed learning curve flattens more than predicted by the dashed power-law fit. (B) ImmSET, evaluated with 180 TCRs per peptide, shows approximately power-law scaling across the same range of training-set sizes. Points denote three sampled datasets; blue and orange traces in (B) denote the ImmSET instance and ensemble models, respectively.

The same nested datasets were used to evaluate the transformer-based ImmSET model. At some smaller data regimes, TACO outperforms individual ImmSET instances reported previously [17]. With larger training sets, those instances overtake TACO. The ImmSET ensemble remains stronger across the reported data range. Thus, TACO’s geometric and physicochemical prior is particularly useful when data are limited, while higher-capacity sequence models derive larger gains from additional data. A direct comparison of these scaling curves is provided in Appendix F.

Overall, TACO extracts useful signal from comparatively modest datasets and remains transparent enough to inspect, but its performance does not continue to improve at the same rate as high-capacity sequence models. Comparisons between model classes should therefore consider performance across training scales, alongside performance at any one dataset size.

## 5 Physicochemical Interpretation

TACO has five amino-acid contact-potential matrices and two peptide-exposure profiles, all released with the model; the complete set is shown in Appendix H. Both exposure profiles assign low weight to peptide position 2 and the C-terminal position, consistent with the B and F anchor residues being buried in the HLA-A*02:01 groove (Figure 3A). Their remaining differences support separate profiles for germline and CDR3 loop groups.

**Fig. 3:**
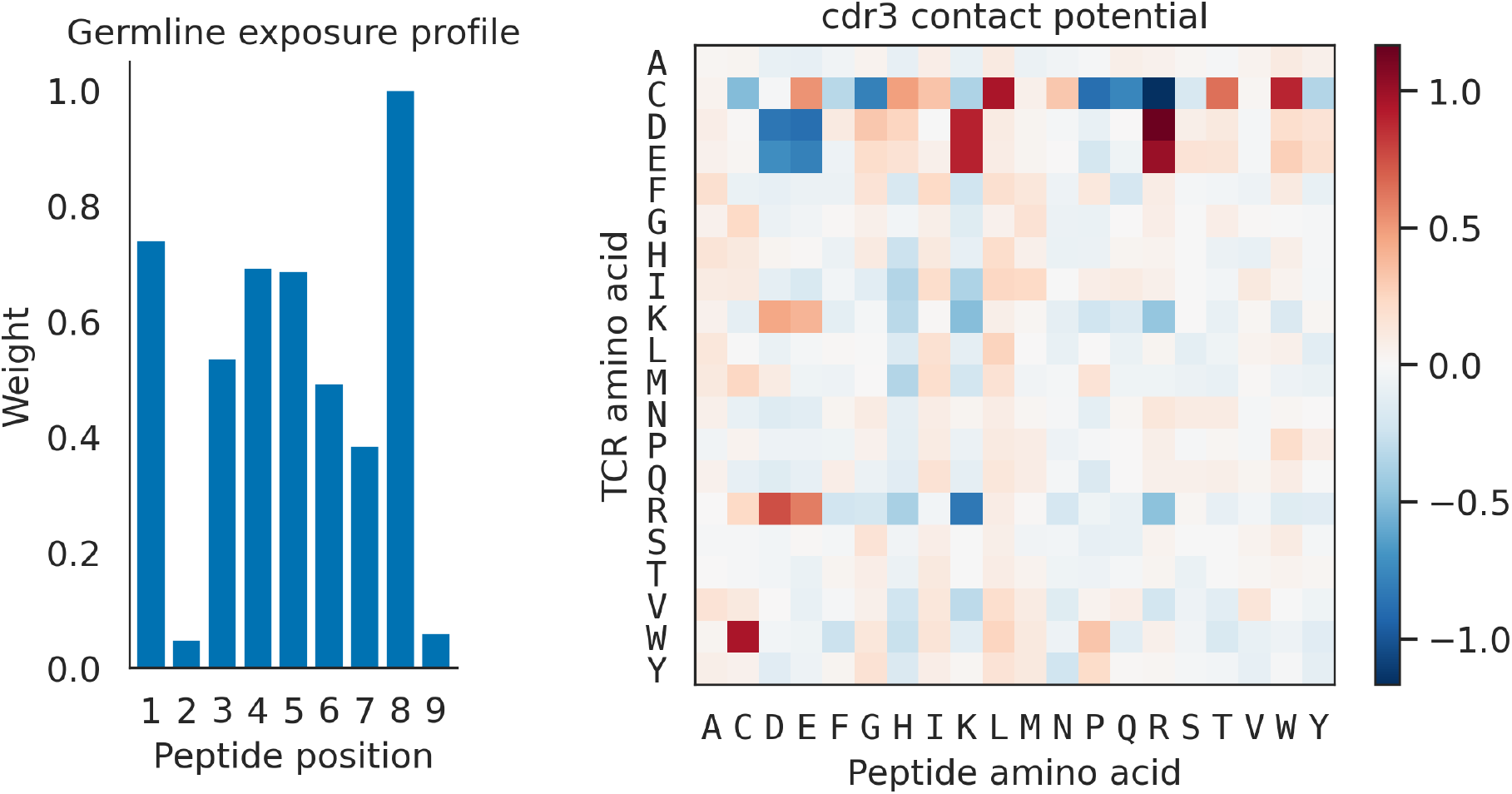
Learned TACO parameters. Left: the germline exposure profile, where near-zero weights at positions 2 and 9 recover the known burial of MHC anchor residues. Right: the CDR3 contact potential, whose dominant pattern is charge selectivity: acidic TCR residues favor basic peptide residues and disfavor acidic peptide residues.

TACO’s parameters are interpretable at the level of loop groups, amino-acid identities, and peptide positions, but they do not constitute a pair-specific TCR–pMHC contact map. The canonical orientation fixes broad, smooth loop footprints over the peptide before model fitting. Thus, a positive contact-potential entry does not imply that a particular CDR residue physically contacts a particular peptide residue in an individual complex. Instead, it describes the learned contribution of a TCR–peptide amino-acid pairing when those residues occur within the corresponding loop-group geometry.

Figure 3 shows the shared CDR3 potential. Its dominant pattern is charge-complementarity: TCR D/E favor peptide K/R and disfavor peptide D/E, while the TCR K/R favor peptide D/E and disfavor peptide K/R. The matrix also contains favorable contributions to cysteine–tryptophan pairings in both orientations (TCR C–peptide W and TCR W–peptide C), nominating sulfur-aromatic interactions as a hypothesis for potential follow up.

The remaining matrices show loop-group-specific patterns. In CDR1 *β*, the nonpolar TCR residue proline shows pronounced charge-selective peptide preferences, favoring residues K and R and disfavoring D and E. CDR2 *β* contains more charge-complementarity effects, including a positive TCR R–peptide E entry paired with a strongly negative TCR R–peptide K entry. In CDR1 *α*, the methionine row exhibits a selective peptide-side-chain signature: lysine, histidine, tyrosine, methionine, and arginine are favorable, whereas glutamate, glutamine, and aspartate are unfavorable. The methionine–tyrosine entry is compatible with a sulfur–aromatic contribution, although the full pattern is broader than aromatic recognition. In CDR2 *α*, tyrosine strongly favors peptide tryptophan and disfavors peptide aspartate. This tryptophan-selective signal is consistent with favorable aromatic packing or *π* interactions, while the aspartate penalty indicates a distinct local chemical constraint. These learned contributions are not direct measurements of physical affinity, but they provide specific hypotheses about residue-pair features associated with activation for structural and experimental follow-up.

## 6 Throughput

TACO performs CPU-only inference using NumPy. We benchmarked it on an Intel Xeon Platinum 8573C CPU with two sockets, 32 physical cores, and 64 logical cores, across workloads ranging from individual TCR–peptide scores to repertoire-scale peptide-panel screening. Multithreaded and single-threaded predictions matched exactly. When evaluating multiple peptides in one run (“panel evaluation”) we can reuse intermediate quantities across peptides and achieve substantially higher per-pair throughput than single-peptide workloads. One million TCRs scored against one peptide reached 17,964 predictions per second on a single thread, while the same repertoire scored against 100 peptides reached 241,267 predictions per second. Multithreading is advantageous at large scale, reaching 406,159 predictions per second for the 100-million-pair panel evaluation, although its overhead makes single-threaded inference faster for small workloads.

These speeds support high-throughput screening of large TCR repertoires against extensive peptide libraries, including viral or proteome-derived candidate panels. Importantly, TACO’s throughput is achieved in the HLA-A*02:01 setting where predictive performance is comparable to AlphaFold3-based prediction; its panel-mode rate is nearly four orders of magnitude greater than that reported for the ImmSET ensemble, a sequence-based deep learning model, and more than eight orders of magnitude greater than those of AlphaFold-derived structural pipelines [17].

## 7 Large-scale viral peptide screening

We next conducted a practical test assessing how TACO’s speed and performance translate to a large screening task using independently-generated data. We focused on the task of nominating candidate antigens for TCRs of unknown specificity but suspected viral association by screening them against large panels of plausible epitopes. For our test, we used publicly-reported and recently-validated TCRs specific for three viral, A*02:01-restricted epitopes: 381 TCRs recognizing GILGFVFTL from influenza A virus, 158 recognizing YLQPRTFLL from SARS-CoV-2, and 91 recognizing NLVPMVATV from cytomegalovirus [15]. These peptides are all dissimilar (a Levenshtein distance of at least four) from all peptides used in TACO’s training and validation, and the TCRs themselves were reported and validated in external work.

For this set of 630 TCRs, we used TACO to score each TCR against a panel of 19,751 predicted HLA-A*02:01-restricted epitopes from reviewed human-virus proteins in Swiss-Prot [1] (Appendix G) and which included all three of the ground truth epitopes for these TCRs. TACO evaluated the resulting 12,443,130 candidate TCR-pMHC combinations in 6.3 minutes. For each TCR, we ranked its annotated target peptide within the overall peptide panel and recorded its rank percentile, where lower values indicate better prioritization and 0.5 is the expected value under random ranking. For individual TCRs across all three epitopes, TACO consistently ranked the correct cognate epitope in the top half of candidates, with the median GIL-specific TCR ranked at the24 *th* percentile, the median YLQ-specific TCR at the 31 *st* percentile, and the median NLV-specific TCR at the25 *th* percentile (Figure 4).

**Fig. 4:**
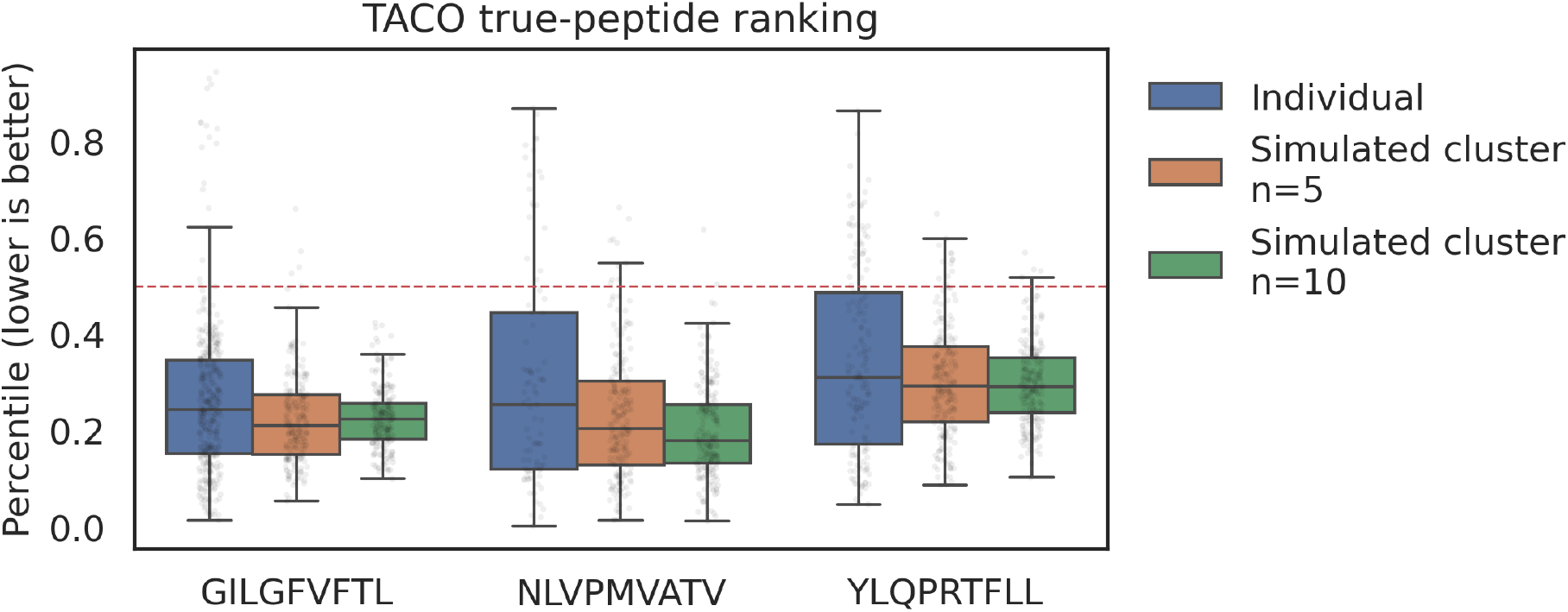
Recovery of known viral targets from a large peptide panel. In 6.3 minutes, TACO ranked the known target peptide for each of 630 TCRvdb TCRs against 19,751 predicted strong HLA-A*02:01-binding viral 9-mers. TCRs were annotated to GILGFVFTL (381 TCRs), NLVPMVATV (91 TCRs), or YLQPRTFLL (158 TCRs). Lower target-rank percentiles indicate better recovery; the dashed line denotes the 50th percentile expected under random ranking. Simulated co-specific groups were formed by averaging scores across 5 or 10 randomly sampled TCRs annotated to the same target peptide, reducing the upper tail of poor target ranks.

We also assessed whether averaging predictions across simulated groups of co-specific TCRs could improve TACO’s ability to give a strong rank to the correct epitope. Here, we averaged peptide scores across groups of *n* = 5 and *n* = 10 randomly-sampled TCRs recognizing the same epitope. For each TCR, we compared its individual target rank to its expected rank when included in a random co-specific cluster; across all three epitopes, clustering significantly improved target ranking (paired Wilcoxon signed-rank test, *p* = 1.2 × 10 ^−5^ for *n* = 5, *p* = 3.3 × 10 ^−8^ for *n* = 10). These groups are simulated rather than inferred biological clonotypes, and testing whether this behavior extends to naturally occurring TCR clusters and broader antigen libraries will be an important direction for future work.

## 8 Discussion

TACO provides an open, compact model for HLA-A*02:01 antigen-induced T-cell activation that avoids perpair structure prediction. Its 2,019 fitted parameters support competitive prediction, CPU-based repertoire-scale screening, and direct inspection of the learned contact potentials and exposure profiles. The TCR*β* -only results also support applications with unpaired repertoire data.

In HLA-A*02:01, TACO achieves performance comparable to AlphaFold3-based prediction while evaluating up to 400,000 TCR–peptide pairs per second in panel mode. This rate is nearly four orders of magnitude greater than that reported for the ImmSET ensemble and more than eight orders of magnitude greater than those of AlphaFold-derived structural pipelines [17], enabling searches against large viral, mutational, or proteome-derived peptide libraries. TACO shows that high-throughput prediction need not depend on making existing complex models faster; it can come from building fundamentally different models informed by the biological and architectural lessons of those models.

More broadly, evaluating TCR–peptide models at a single training scale cannot establish their long-term potential. TACO learns efficiently in limited-data regimes, but its gains flatten with additional training data, whereas higher-capacity sequence models continue to improve. Future structured extensions could incorporate HLA context, richer positional interactions, and paired-chain information while retaining interpretable components. Learning curves across data scales will be essential for comparing these approaches. Such studies should also test whether sequence- and structure-based models make complementary errors, rather than treating either as a single global baseline. TACO is specific to HLA-A*02:01; extension to other alleles will require allele-specific data and likely allele-specific contact potentials, exposure profiles, and geometry.

The released code and parameters provide an open baseline for these tests and for broader exploration of fast, interpretable TCR–peptide prediction. A model need not perfectly resolve every TCR–peptide pair to be useful: at the scale enabled by TACO, its rankings can focus experimental effort on more promising candidates, as illustrated here by prioritization of candidate peptides for experimental follow-up. We hope the community will use this capability to build and test workflows that have not previously been practical, establishing the experimental and computational infrastructure needed to take advantage of future models that retain this throughput while improving predictive performance.

## Acknowledgments

We thank Phil Bradley for sharing the AF3-TCRdock evaluation result reported in [17]. The authors used Claude 4.6 Opus to assist with development and iteration of research code, including code used to generate schematic figures. All AI outputs were reviewed for accuracy by the authors.

## Data and Code Availability

The TACO scorer class, trained model parameters (JSON), training and evaluation notebooks are available at https://github.com/mgarcianoceda/TACO. This study uses proprietary MIRA [11] and pairSEQ [8] TCR–pMHC interaction data: amino acid sequences paired with binary CD8+ T-cell activation labels. The training data are owned by Adaptive Biotechnologies and are not publicly available. Related datasets are available for licensed use through the Harell Data platform (https://harelldata.com/). For results obtained on the IMMREP25 benchmark [16], and results obtained using TCRvdb [15] and Swiss-Prot [1], those datasets are publicly available.

