## Appendices for "TACO: An open, interpretable model for microsecond-scale TCR specificity prediction"

#### Appendix A Retrospective Model-Variant Comparisons

We evaluated a set of model variants on the primary held-out set. All values in Table A1 use the same held-out examples; per-peptide AUROC is the primary metric, and overall AUROC is shown for reference.

The comparisons support several design choices in TACO. The validation-selected orientation and fitted exposure profiles each improve holdout performance over their structural or hand-specified counterparts. Retaining separate loop groups is important: performance decreases as loop classes are pooled, and a single shared potential retains little of the full-model signal. Splitting the shared CDR3 group into separate CDR3$\alpha$ and CDR3$\beta$ potentials does not improve performance. The two-profile model provides a modest improvement over a single fitted shared profile. Geometry and peptide orientation are also important: removing the distance dependence of the geometry, inverting the anchor profile, or reversing the profile along the peptide substantially reduces performance.

Table A1: Comparison of TACO model variants on the primary held-out set. Per-peptide AUROC is the primary evaluation metric.

| Variant | Per-peptide AUROC | Overall AUROC |
|---|---|---|
| **Final model: fitted profiles, selected orientation** | **0.6661** | **0.6659** |
| Fitted profiles, structural orientation | 0.6535 | 0.6545 |
| Symmetric contact potentials | 0.6543 | 0.6567 |
| Hand-specified profiles, selected orientation | 0.6561 | 0.6576 |
| Hand-specified profiles, structural orientation | 0.6442 | 0.6470 |
| Six surfaces: one potential per loop | 0.6568 | 0.6575 |
| Three surfaces: CDR1, CDR2, and CDR3 pooled across chains | 0.6429 | 0.6442 |
| One surface: all loops share one potential | 0.6017 | 0.5989 |
| One surface, strict orientation, prior profile | 0.5870 | 0.5831 |
| Charge-only, strict orientation, prior profile | 0.5684 | 0.5690 |
| Geometry removed: $\sigma \to \infty$ | 0.5809 | 0.5771 |
| One fitted shared exposure profile | 0.6634 | 0.6663 |
| Inverted-anchor exposure profile | 0.6419 | 0.6461 |
| Reversed exposure profile | 0.6066 | 0.6019 |

Figure A1 compares representative variants as the number of training peptides increases, with 180 TCRs retained per peptide. The five-surface model improves substantially over the one-surface and charge-only variants at all dataset sizes. Fitting the exposure profiles gives an additional improvement that is most evident in the larger subsets. These comparisons show that the full model's performance depends on the combination of residue-level contact potentials, loop-specific surfaces, peptide-position weighting, and the canonical orientation.

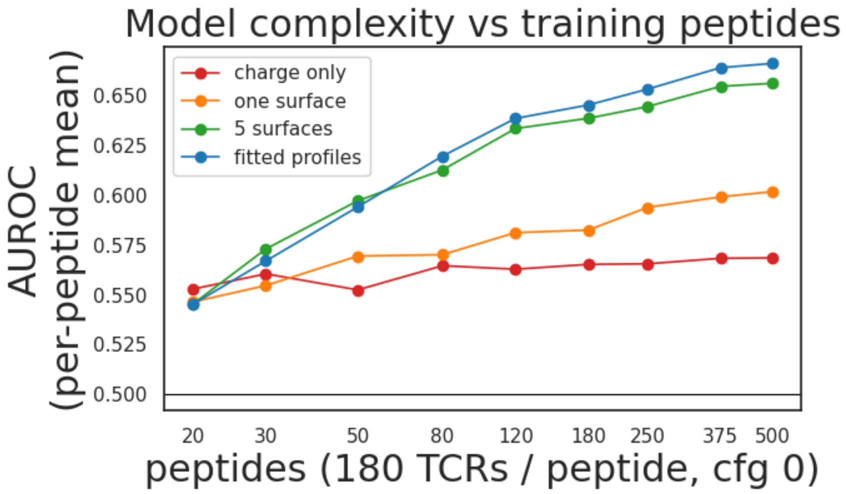


Fig. A1: Scaling of representative TACO variants with the number of training peptides. Each point reports mean per-peptide AUROC on the held-out set, using 180 TCRs per training peptide from Dataset 0. The full model with fitted exposure profiles outperforms the simpler charge-only and one-surface variants across the evaluated range.

#### Appendix B Orientation selection

Table B1 gives the seven hand-specified orientations evaluated during model selection. Each row defines a complete geometry hypothesis. For every candidate, contact potentials and peptide-exposure profiles were fitted independently using the same training data, and validation AUROC was used to select the final orientation.

SP scales the displacement of each germline-loop center from peptide position 5, with SP= 1 retaining the hand-specified centers. GD places the CDR3$\alpha$ and CDR3$\beta$ centers at $\mu = 5 - \text{GD}$ and $\mu = 5 + \text{GD}$, respectively. SIG_G and SIG_C specify the contact widths $\sigma$ for germline and CDR3 loops. SW scales the hand-specified sweep slopes $\rho$; SW= 0 sets all sweep slopes to zero.

Table B1: Candidate orientations used for orientation selection.

| Candidate | SP | GD | SIG_G | SIG_C | SW | Validation AUROC |
|---|---|---|---|---|---|---|
| Structural guess | 1.0 | 1.5 | 1.5 | 1.5 | 1.0 | 0.6846 |
| No sweep | 1.0 | 1.5 | 1.5 | 1.5 | 0.0 | 0.6973 |
| Wider contacts | 1.0 | 1.5 | 2.0 | 2.0 | 0.0 | 0.6997 |
| Wider CDR3 only | 1.0 | 1.5 | 2.0 | 2.6 | 0.0 | 0.6997 |
| **CDR3 pushed apart** | **1.0** | **2.0** | **2.0** | **2.6** | **0.0** | **0.6999** |
| Germline spread 1.4 | 1.4 | 2.0 | 2.0 | 2.6 | 0.0 | 0.6993 |
| Germline spread 1.8 | 1.8 | 2.0 | 2.0 | 2.6 | 0.0 | 0.6897 |

### Appendix C Fitting details

#### Appendix C.1 Objective

For a fixed candidate orientation, TACO learns the five contact-potential matrices $M = \{M^{(r)}\}_{r=1}^{5}$, two peptide-exposure profiles, and global intercept $\beta_0$ by minimizing penalized binary cross-entropy over the $N$ observed training rows:

$$\mathcal{L} = \frac{1}{N}\sum_{n=1}^{N}\left[\log(1+\exp(z_n+\beta_0)) - y_n(z_n+\beta_0)\right] + \lambda\|\operatorname{vec}(M)\|_2^2, \quad (8)$$

where $y_n \in \{0, 1\}$ is the activation label for row $n$, $z_n$ is the contact score in Equation 6 before addition of the intercept, and $\lambda = 10^{-6}$. The ridge penalty applies only to the contact-potential entries.

The exposure profiles are parameterized by nine control points for the germline loop groups and nine control points for the CDR3 loop group. These control points are interpolated to the length of each peptide before evaluating Equation 6.

#### Appendix C.2 Alternating optimization

The joint objective is not globally convex because each contact contribution combines an exposure weight with a contact-potential entry. We therefore fit the model by alternating between two conditional updates. With the exposure-profile control points fixed, we optimize the five contact-potential matrices and global intercept. With the contact potentials fixed, we optimize the two exposure profiles through their control points.

Each conditional update has a convex objective: with exposure profiles fixed, the score is affine in the contact potentials and intercept; with contact potentials fixed, it is affine in the exposure-profile control points. We use L-BFGS for both updates and repeat the alternating procedure is done for three alternating rounds; each conditional sub-problem is solved by L-BFGS to a gradient tolerance of $10^{-9}$. The iterate with the highest validation AUROC across rounds is retained.

For every candidate orientation, fitting begins from zero contact potentials and the marginal log-odds as intercept. Two profile initializations are tried in parallel — the structural prior and a flat (uniform) profile — and the one yielding the higher validation AUROC is kept. The fit proceeds in two stages: first a single shared potential and profile to obtain a warm start, then five independent potentials with two exposure-profile classes (germline and CDR3), again tried from both the warm start and the prior. Contact potentials, exposure profiles, and intercept are fitted independently for each candidate orientation before validation AUROC is used to select the final orientation.

#### Appendix C.3 Implementation

All optimization uses the exact full-batch loss and gradient over the observed training rows, without class rebalancing or negative subsampling. The implementation avoids explicitly constructing the full $N \times 2{,}000$ design matrix. Instead, rows are processed in blocks grouped by peptide length and antigen chunk. Within each block, all TCRs are scored against all peptides through matrix multiplication, and the corresponding transpose operation yields the gradient contribution for the contact potentials.

Training uses a single NVIDIA T4 GPU, with linear-algebra operations implemented in PyTorch. The released code contains the exact optimizer settings, interpolation procedure, initialization, and convergence criteria.

### Appendix D Structural models predictions

*AlphaFold3-TCRdock:* The AlphaFold3-TCRdock AUROC reported in this study is reproduced from Ref. [17]. The reported metric is for a TCRDock-style adaptation of AlphaFold3 for TCR–pMHC modeling. The adaptation incorporated inter-chain template features and TCR–pMHC docking templates sampled from canonical binding modes of known complexes, extending the template-based strategy introduced for AlphaFold2 in TCRDock [4].

*OpenFold3 preview 2 (OF3p2):* We ran OpenFold3 'preview 2' [22, 2] on the full IMMREP25 A*02:01 dataset (5,000 rows and 10 peptides) using a single seed and diffusion sample. We precomputed MSAs and template matches using the immune-specific search datasets from tcrmodel2 [25]. We predicted TCR-pMHC activation for OpenFold3 using a custom ipTM confidence score for the TCR-pMHC interface.

*OpenDDE:* We ran OpenDDE [3] v1 parameters on the full IMMREP25 A*02:01 dataset (5,000 rows and 10 peptides) using a single seed and diffusion sample. We precomputed MSAs and template matches using the immune-specific search method and datasets from tcrmodel2 [25].

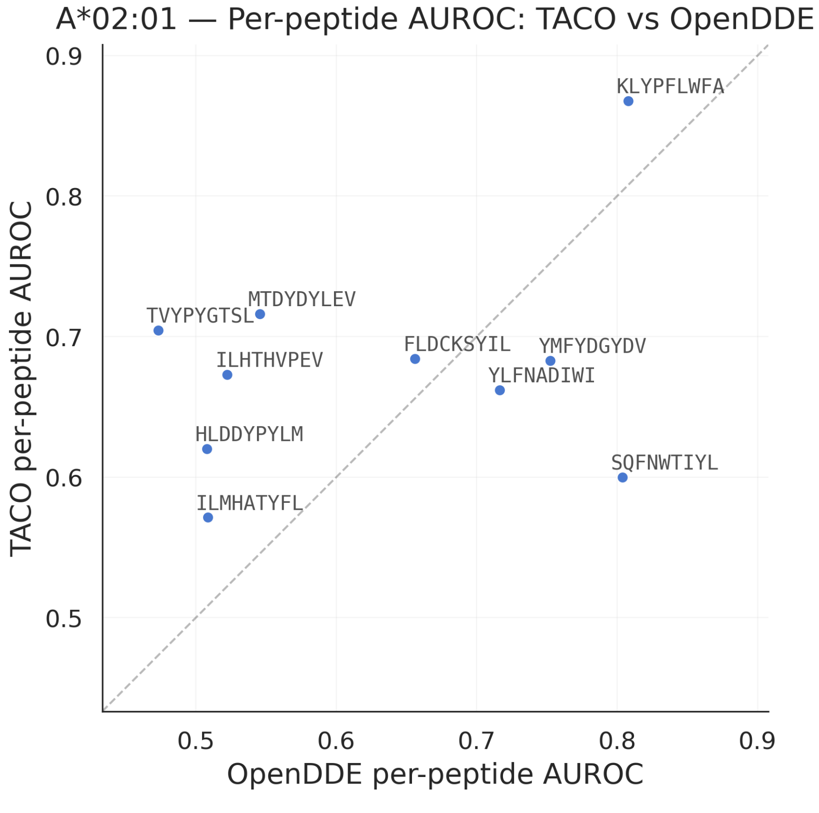


Fig. D1: Per-peptide AUROC of TACO versus OpenDDE on IMMREP25 HLA-A*02:01. Each point is a peptide; distance from the diagonal reflects peptide-specific differences between models.

TACO and OpenDDE show different patterns of per-peptide performance on IMMREP25 HLA-A*02:01. Some peptides are ranked well by TACO and poorly by OpenDDE, while others show the opposite pattern. Figure D1 shows this relationship for the open structural model with the strongest performance in our comparison. The pattern suggests that sequence and structural models may capture partially distinct signal and motivates future evaluation of ensemble predictions.

#### Appendix E Cross-HLA generalization

Despite being trained exclusively with A*02:01 data, TACO generalizes to other HLAs. Although TACO generalizes to other HLAs, its predictive signal is weaker than that observed for A*02:01 (see Figure E1).

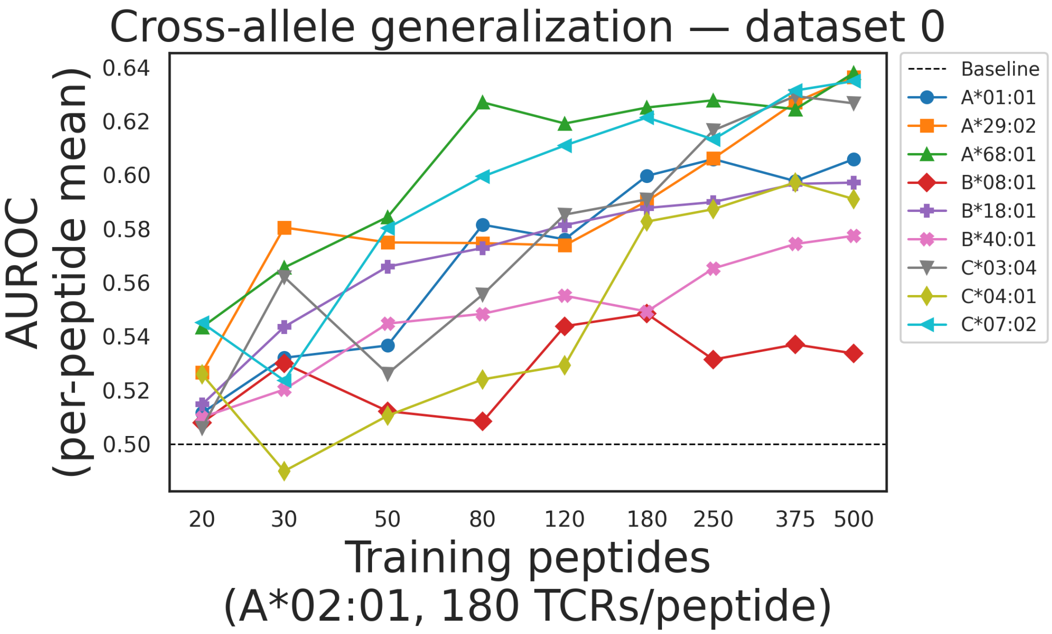


Fig. E1: TACO generalizes to other alleles despite being trained only with A*02:01 data

#### Appendix F Full TACO scaling results

Figure F1 shows TACO scaling with peptides across all TCRs-per-peptide settings, showing it does not follow a power law.

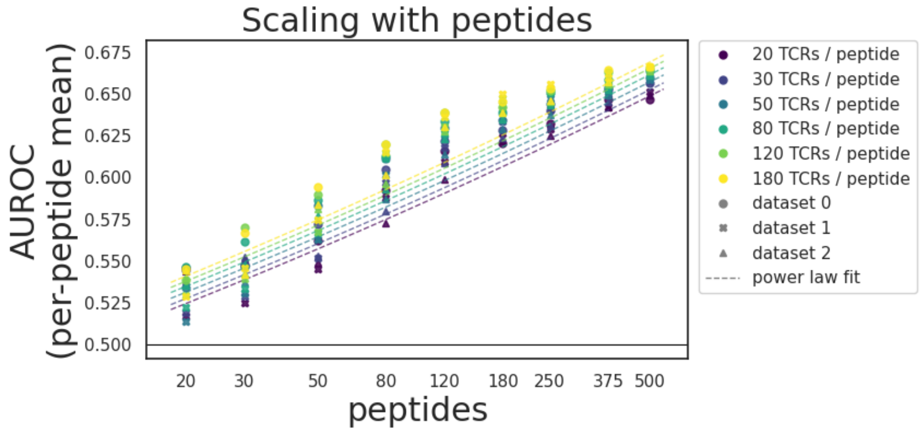


Fig. F1: Full scaling trajectories for TACO.

#### Appendix G Viral peptide screening

We assembled the viral peptide search set used in Section 7 as follows. We first downloaded FASTA files from Swiss-Prot on October 24, 2026, using the query `(virus_host_id:9606) AND (reviewed:true)`. From these files, we generated 1,356,628 9-mer peptides by running a sliding window across all protein sequences. From these candidate peptides, we retained 19,751 predicted HLA-A*02:01 epitopes by selecting peptides that were predicted "strong binders" (`el_rank_percentile` below 0.5%) by NetMHCpan-4.1 [18]. Restricting the search set to predicted A*02:01 epitopes (rather than all 9-mer peptides) allows us to measure the additional information TACO provides at the TCR specificity level, orthogonal to the strong restriction placed by peptide-MHC compatibility that is separately predicted by well-established tools like NetMHCpan-4.1. All three of the ground truth peptides for the TCRs in this experiment (GILGFVFTL from influenza A virus, YLQPRTFLL from SARS-CoV-2, and NLVPMVATV from cytomegalovirus) are present in this final search set of 19,751 peptides.

#### Appendix H Learned Parameters

We can visualize and do some analysis for the two learned peptide-exposure profiles and the five learned contact-potential matrices. In every contact-potential plot, rows denote TCR amino-acid types and columns denote peptide amino-acid types; positive values increase the model score and negative values decrease it. Color ranges differ between panels, so numerical labels should be used when comparing values across surfaces.

##### Appendix H.1 Peptide-exposure profiles

Figure H1 shows the two exposure profiles, each represented by nine control points for a 9-mer peptide. Both profiles assign little weight to position 2 and the C-terminal position, consistent with the B and F anchors occupying HLA-A*02:01 binding pockets. The profiles differ across the remaining positions, supporting the use of separate germline and CDR3 exposure profiles.

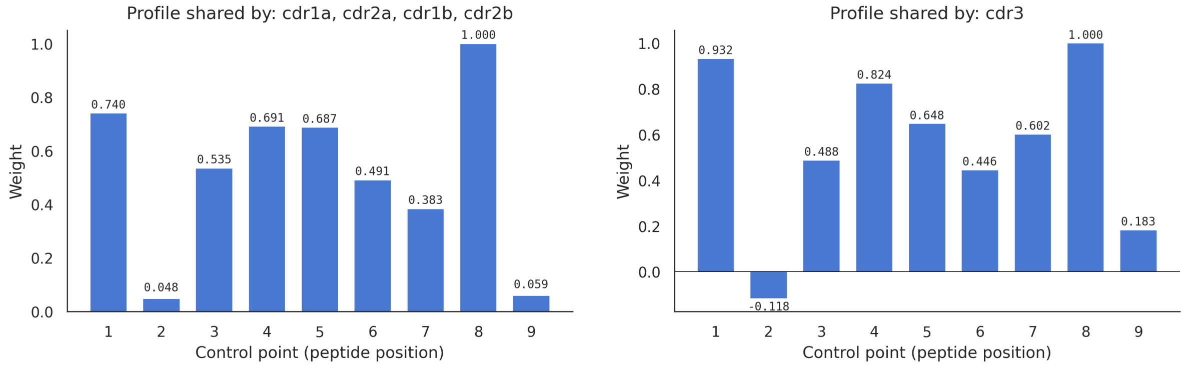


Fig. H1: Learned peptide-exposure profiles. **Left:** profile shared by CDR1$\alpha$, CDR2$\alpha$, CDR1$\beta$, and CDR2$\beta$. **Right:** profile for the shared CDR3$\alpha$/CDR3$\beta$ surface. Values are interpolated between control points for peptides whose length differs from nine residues.

#### Appendix H.2 Contact potentials

The contact potentials show distinct patterns across surfaces. The shared CDR3 potential is dominated by charge-selective interactions: TCR D/E favour peptide K/R and disfavor peptide D/E, with corresponding structure in the K/R rows. It also contains positive cysteine–aromatic entries, including C–W and W–C. CDR1$\beta$ has the largest individual coefficients, notably P–K (+2.88) and P–E (−2.61), indicating strong charge-selective behaviour from an uncharged TCR residue. CDR2$\beta$ distinguishes peptide D and E in several rows, including R–D (+2.14) and R–E (−1.52). CDR2$\alpha$ contains prominent tyrosine-associated entries, including Y–D (−1.96) and Y–W (+1.15). The CDR1$\alpha$ potential is more heterogeneous, with no similarly dominant single residue class.

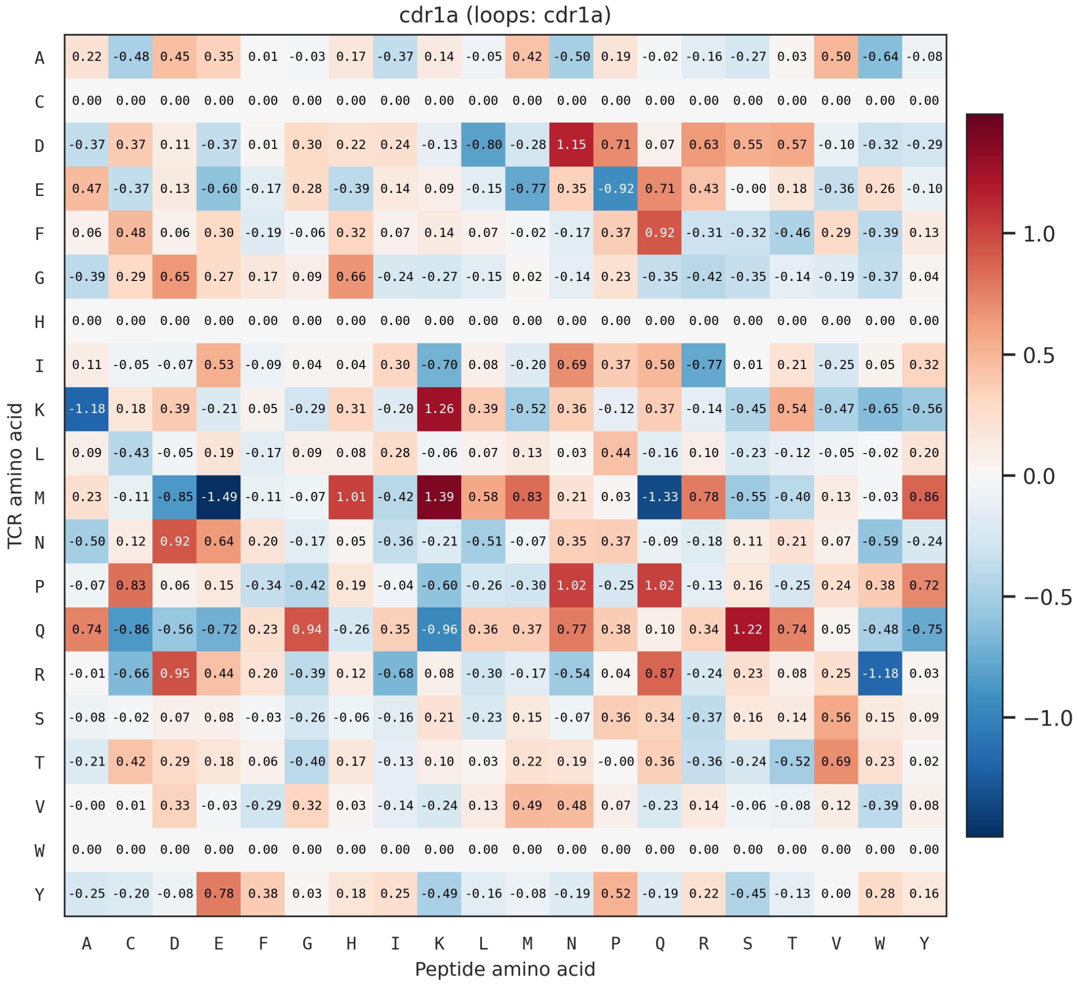


Fig. H2: Learned contact-potential matrix for the CDR1$\alpha$ loop. The TCR methionine row exhibits a selective pattern, favoring peptide K (+1.39), H (+1.01), Y (+0.86), M (+0.83), and R (+0.78), while disfavoring E (−1.49), Q (−1.33), and D (−0.85).

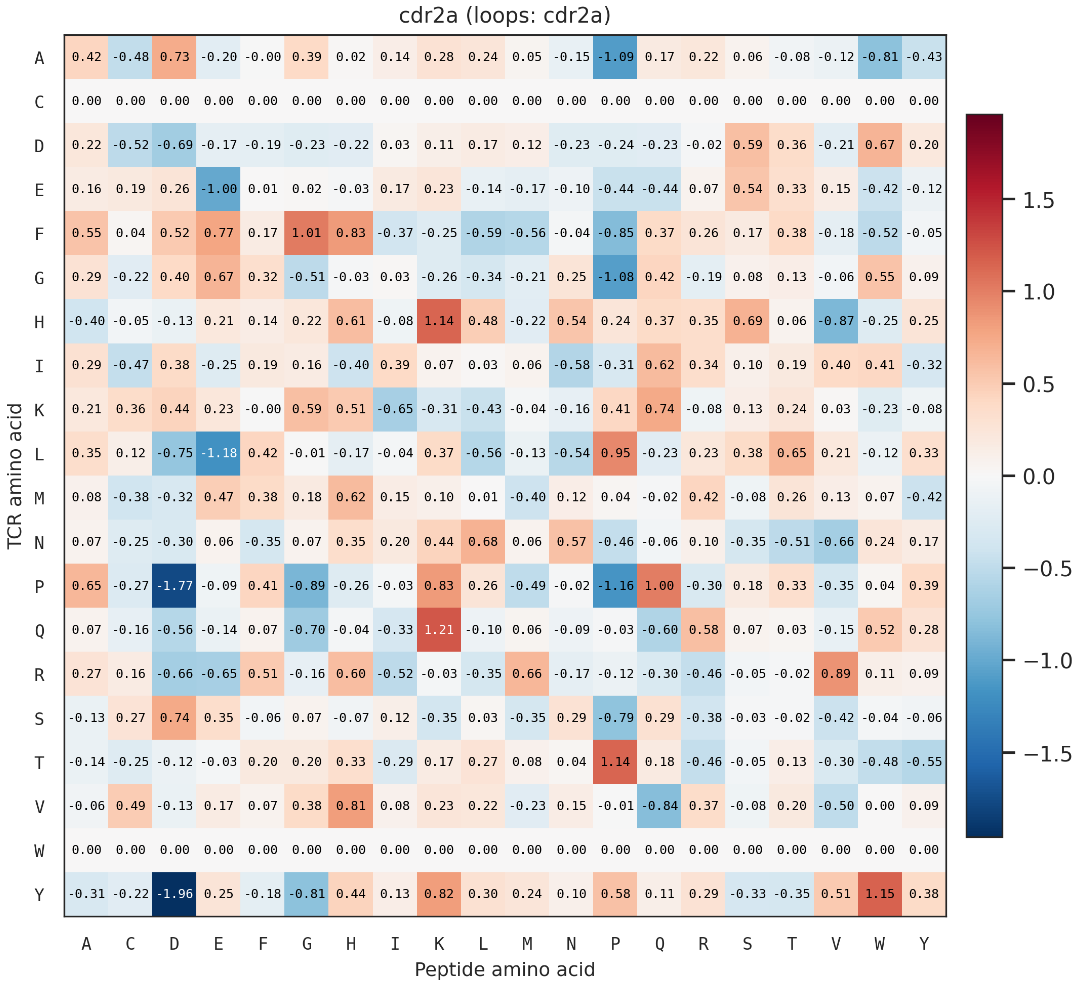


Fig. H3: Learned contact-potential matrix for the CDR2$\alpha$ loop. The TCR tyrosine row strongly favors peptide W (+1.15) and disfavors peptide D (−1.96). The favorable Tyr–Trp entry nominates aromatic packing as a structural hypothesis for follow-up.

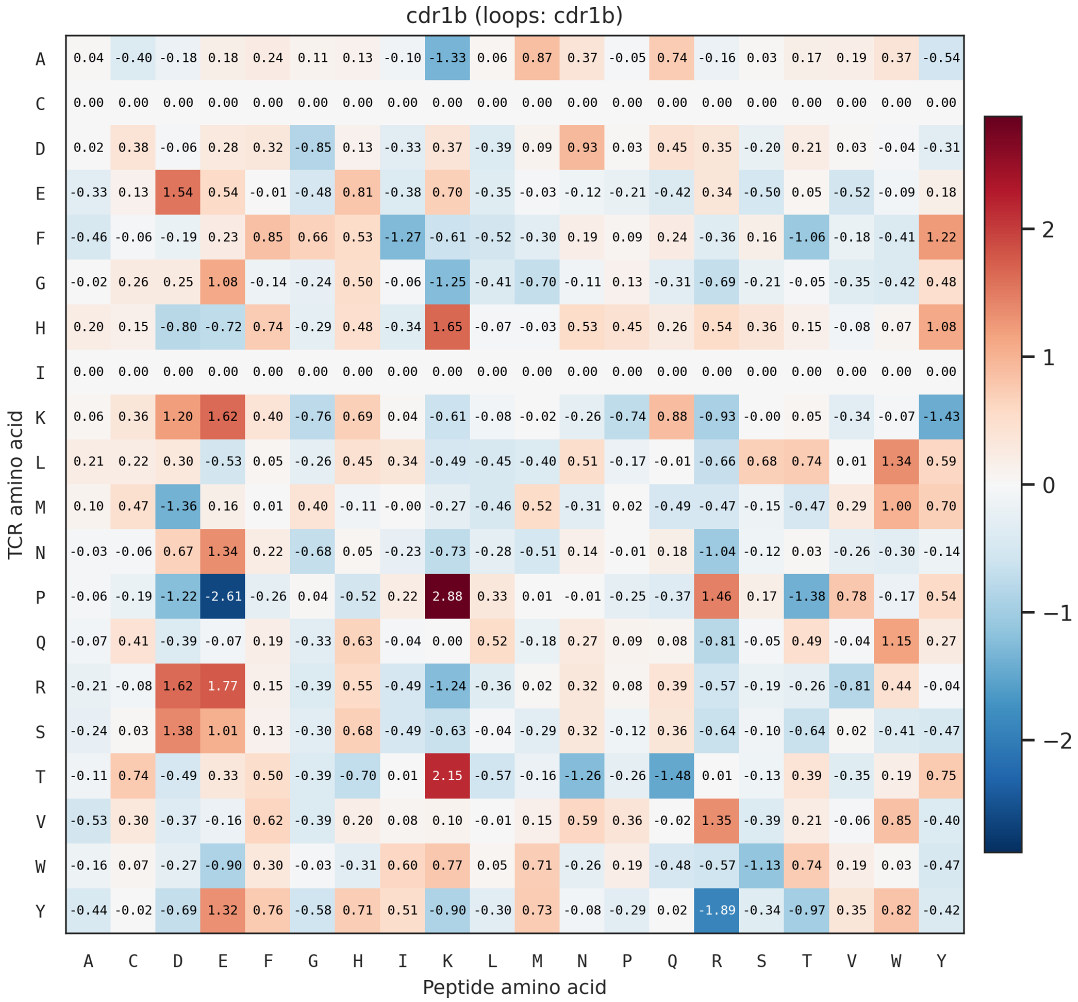


Fig. H4: Learned contact potential for the CDR1$\beta$ loop. The largest coefficients are P–K (+2.88) and P–E (−2.61), indicating charge-selective effects from proline in this fitted potential.

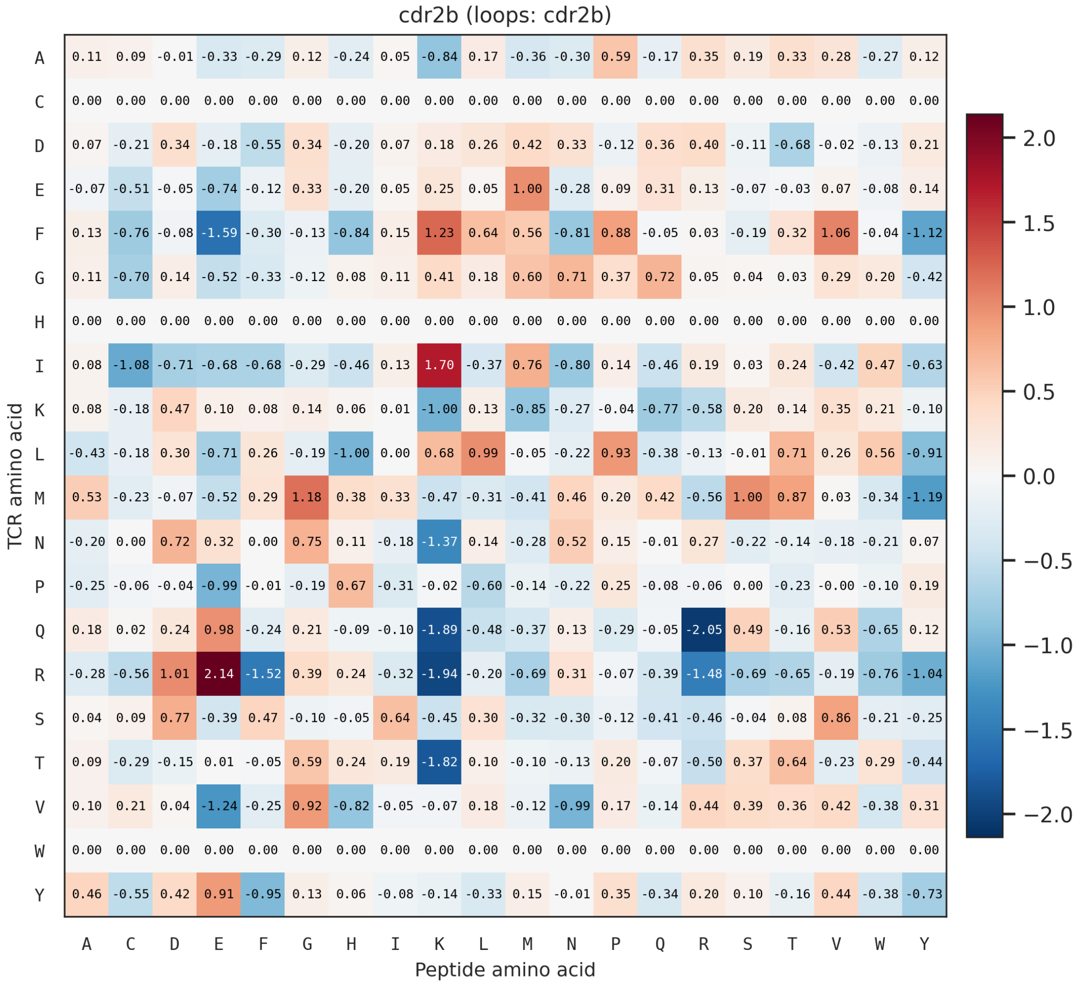


Fig. H5: Learned contact-potential matrix for the CDR2$\beta$ loop. The TCR arginine row shows a marked charge-complementary preference, favoring peptide glutamate (+2.14) while strongly disfavoring peptide lysine (−1.94).

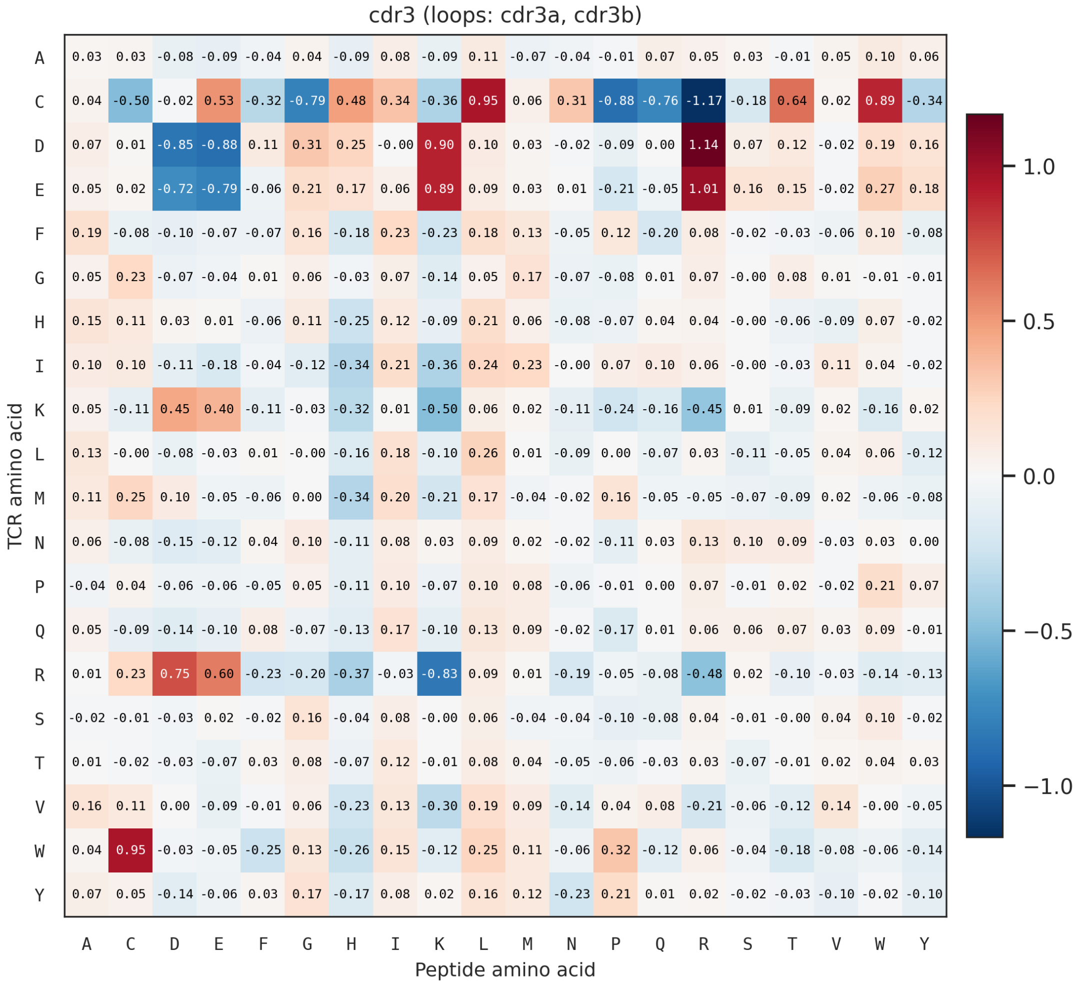


Fig. H6: Learned contact potential shared by the CDR3$\alpha$ and CDR3$\beta$ loops. Charged TCR residues show a coherent charge-selective pattern, and positive cysteine–aromatic entries include C–W and W–C.
